# A New Paradigm for Applying Deep Learning to Protein-Ligand Interaction Prediction

**DOI:** 10.1101/2023.11.01.565115

**Authors:** Zechen Wang, Sheng Wang, Yangyang Li, Jingjing Guo, Yanjie Wei, Yuguang Mu, Liangzhen Zheng, Weifeng Li

## Abstract

Protein-ligand interaction prediction poses a significant challenge in the field of drug design. Numerous machine learning and deep learning models have been developed to identify the most accurate docking poses of ligands and active compounds against specific targets. However, the current models often suffer from inadequate accuracy and lack practical physical significance in their scoring systems. In this research paper, we introduce IGModel, a novel approach that leverages the geometric information of protein-ligand complexes as input for predicting the root mean square deviation (RMSD) of docking poses and the binding strength (the negative value of the logrithm of binding affinity, pKd) with the same prediction framework. By incorporating the geometric information, IGModel ensures that its scores carry intuitive meaning. The performance of IGModel has been extensively evaluated on various docking power test sets, including the CASF-2016 benchmark, PDBbind-CrossDocked-Core, and DISCO set, consistently achieving state-of-theart accuracies. Furthermore, we assess IGModel’s generalization ability and robustness by evaluating it on unbiased test sets and sets containing target structures generated by AlphaFold2. The exceptional performance of IGModel on these sets demonstrates its efficacy. Additionally, we visualize the latent space of protein-ligand interactions encoded by IGModel and conduct interpretability analysis, providing valuable insights. This study presents a novel framework for deep learning-based prediction of protein-ligand interactions, contributing to the advancement of this field.

Key Messages

- We introduce the first framework for simultaneously predicting the RMSD of the ligand docking pose and its binding strength to the target.
- IGModel can effectively improve the accuracy of identifying the near-native binding poses of the ligands, and can still outperform most baseline models in scoring power, ranking power and screening power tasks.
- IGModel is still ahead of other state-of-the-art models in the unbiased data set and the target structure predicted by AlphaFold2, proving its excellent generalization ability.
- Latent space provided by IGModel learns the physical interactions, thus indicating the robustness of the model.

## Introduction

Understanding the precise binding poses of ligands within protein receptor structures holds immense significance in the field of drug design[1, 2, 3, 4]. By accurately predicting ligand binding poses, researchers gain valuable insights into the intricate mechanisms underlying drug-target interactions. This knowledge empowers them to optimize binding affinity, selectivity, and pharmacological properties, ultimately leading to the development of more potent and efficient therapeutic agents[5, 6].

Although experimental methods can be employed to determine the binding mode of small molecules within proteins, their efficiency tends to be relatively low, and their accuracy is not always guaranteed[7, 8]. Experimental technique such as X-ray diffraction offers atomic insights into proteinligand interactions, is often time-consuming, resource-intensive, and may encounter challenges in obtaining high-resolution structures or capturing dynamic aspects of the binding process[9, 10]. Additionally, experimental approaches may be limited by the availability of suitable protein samples and the complexity of the system under investigation. Consequently, relying solely on experimental techniques for ligand binding pose determination may not always meet the demands of high-throughput drug discovery and design[11].

Computational methods, such as molecular docking, play a pivotal role in this endeavor, offering efficient and accurate tools to explore the vast chemical space and guide rational drug design efforts[12, 13, 14, 15]. To guide fast and accurate protein-ligand complex structure prediction, especially the ligand binding pattern (given that the protein structure is already known or determined), the scoring functions (SFs) are developed[16, 17]. SFs are used to calculate the binding energy or score associated with each pose, allowing for the ranking and selection of the most favorable binding configurations[17, 9]. During ligand pose prediction, SFs consider various factors, such as the complementarity of molecular shapes, electrostatic interactions, van der Waals forces, hydrogen bonding, and solvation effects[11]. By evaluating these energy terms, SFs estimate the overall binding affinity or likelihood of a ligand pose being biologically relevant. SFs help guide the exploration of ligand conformational space, identifying potential binding modes and providing insights into the binding strength and specificity of ligand-receptor interactions[18]. They enable researchers to prioritize ligand poses for further analysis or optimization, facilitating the rational design of drug candidates with improved binding affinity and selectivity[9, 19].

In the 1980s, people began to design rules for evaluating ligand binding mode based on physical knowledge and expert experience, or using the force field parameters from bio-molecule simulation systems, which was the early SF[20]. The traditional SF is usually a polynomial with few parameters and atom type definitions, thus is not accurate enough to describe the conformational space of the protein-ligand complex[21]. Another category of SF involves machine learning (ML)-based approaches, which have been traced back to 2004[22]. This kind of SF predicts the ligand binding affinity by extracting interaction features using various machine learning techniques[23, 24]. Over the years, numerous notable SFs have been proposed, including RF-Score[25] (as well its virtual screen version, RF-score-VS), NN-Score[26], AGL-Score[27], etc. These SFs have achieved excellent performance (“scoring power”[28]) under the PDBbind related benchmarks. The growing availability of public experimental data and advancements in DL algorithms have led to a rising trend of utilizing DL models for predicting protein-ligand interactions[29, 30, 31]. Since 2017, numerous DL-based models have been introduced for predicting protein-ligand affinity. For example, AtomNet[32], K_*deep*_[33] and Pafnucy[34] employed 3D convolutional neural networks (CNN) to establish a connection between the 3D mesh representation of the complex and its affinity. Additionally, our group has contributed two 2D convolution-based models, namely OnionNet and OnionNet-2, which assess protein-ligand affinity by counting the number of contacts between ligand atoms and protein atoms/residues in different distance intervals[35, 36]. However, recent studies have revealed that ML or DL models trained with only native structures exhibit limited performance when applied to docking power and screening power tasks, despite their strong performance in scoring power[37]. In order to make the model applicable to practical scenarios, researchers began to design predictors that can be directly applied to identify near-native poses of the ligand or screening active compounds. For instance, DeepBSP employed 3D CNN to predict the RMSD of ligand docking poses[38].The DeepRMSD+Vina, previously proposed by our group, adopts the modified formats of van del waals and columbic terms used in many traditional force fields in molecular mechanics as features and has been demonstrated to be effective in docking power and docking pose optimization[39]. In addition, DeepDock utilized a graph neural network to learn the distance probability distribution between protein-ligand atoms, rather than directly predicting the protein-ligand binding affinity or the ligand pose RMSD[40]. However, it turns out that many DL scoring functions could not perform well for all evaluation metrics[28]. Building upon the idea of DeepDock, RTMScore introduced a novel graph representation and employed a graph transformer model to learn distance probability distributions, achieving state-of-the-art results across multiple virtual screening test sets and docking poses prediction tasks[41]. More researches now realized the importance to have balanced performance for both docking and screening tasks[42, 43, 44].

Among the numerous deep learning methods mentioned above, they either provide affinity prediction or ligand pose prediction, with few being able to simultaneously offer directly interpretable indicators with physical meanings, which is more intuitive for computational drug developers[45, 28]. Other methods lack robustness, especially when using predicted structures or cross-docked structures as protein templates, there are few approaches that can accurately predict the optimal ligand binding poses[46, 47].

To these ends, we proposed a SF based on the geometric graph neural network, named IGModel, to further elevate the upper limit of protein-ligand interaction prediction, especially the docking poses prediction. The input of IGModel includes two graphs: one is the protein binding pocket graph, and the other is the protein-ligand atomic interaction graph. Unlike previous graph representations[48, 49], IGModel integrates the distance and orientation between interacting atoms as geometric features of the complex to provide a more comprehensive description of the relative positions of the ligand within the binding pocket. We employ EdgeGAT layers proposed by Kamiński et al.[50] to encode the protein-ligand interaction to obtain a latent vector which is further decoded into the RMSD of the pose and the binding strength to the protein through two decoding modules. Therefore, IGModel can be subdivided into two sub-branches, namely IGModel_*RMSD*_ and IGModel_*pkd*_. In the CASF-2016[28] docking power test, IGModel_*RMSD*_ achieved the highest Top1 docking success rate (97.5% and 95.1% when including and excluding the native poses, respectively). On cross-docking datasets like PDBbind-CrossDocked-Core set[51] and DISCO set[52], IGModel still performs well, which is comparable or even better than other baseline models. In addition, IGModel can also show excellent performance on the unbiased test set and the datasets containing target structures generated by AlphaFold2[53], proving that IGModel has outstanding generalization ability and are practical for drug discovery. The model captures key charge-charge or hydrogen bond interactions indicating potential robustness for protein-ligand interaction prediction and the model thus could also be used for lead optimization. Overall, we present a highly accurate ligand pose and binding affinity prediction model which would be an useful tool for drug design.

## Methods

### Datasets

In this study, the native protein-ligand complexes from the PDBbind database (v.2019)[54] along with their corresponding docking poses are utilized for training. In order to increase the diversity of ligand binding conformations, an average of 15 docking poses are generated by AutoDock Vina[55] and ledock[56] for each native protein-ligand complex. Samples from CASF-2016 and those with peptide ligands were excluded, as well as any samples that failed to be parsed by certain programs, such as rdkit[57]. The actual root mean square deviation (RMSD) of docking poses are calculated using the spyrmsd package[58]. The binding affinity (pKd) of the native protein-ligand complex is represented as the negative logarithm of the dissociation constant (K_*d*_), inhibition constant (K_*i*_), and half-inhibitory concentration (IC50). The parameters for molecular docking and the distributions of RMSD and pKd of docking poses are shown in Supplementary part 1.

### Graphical representation of protein-ligand complexes

Physically, ligand binding within the pocket is influenced by non-bonding interactions, and the potential energy surfaces of protein-ligand binding reveal geometric preferences among atoms. Therefore, accurately describing the relative positions of atoms in the protein-ligand system is crucial for characterizing the protein-ligand interactions. In this study, we construct a heterogeneous graph G^*RR*,*LL*,*RL*^ to encode protein-ligand interactions. This graph comprises two node types: protein atom nodes (V^*R*^) and ligand atom nodes (V^*L*^). To conserve computational resources, only protein atoms within 8 Å*A* of the ligand co-crystal structure are used for interaction modeling, without considering all atoms. There are four message-passing channels in G^*RR*,*LL*,*RL*^, namely E^*RR*^ (V^*R*^ to V^*R*^), E^*LL*^ (V^*L*^ to V^*L*^), E^*RL*^ (V^*R*^ to V^*L*^), and E^*LR*^ (V^*L*^ to V^*R*^).

Within the ligand subgraph G^*L*^=(V^*L*^,E^*LL*^), we define seven types of ligand atoms: C, N, O, P, S, Hal (representing halogen elements F, Cl, Br, and I), and DU (representing other element types). If a covalent bond exists between atoms i and j, an edge e_*ij*_ ^*L*^ is defined between nodes v_*i*_^*L*^ and v_*j*_ ^*L*^. The rdkit package enables the extraction of physical and chemical information related to the chemical bond, which is used as the edge feature in conjunction with the bond length. For the protein subgraph G^*R*^=(V^*R*^,E^*RR*^), we redefined atom types based on the element type, the residue type it blongs to, and whether it is located in the main chain or the side chain. For example, “LYS-MC” and “LYS-SC” represent the C atom on the main chain and side chain of LYS, respectively. The one-hot encoding of atom types, along with aromaticity, charge, and the distance to the *α*-C atom, are collectively used as node features for G^*R*^. When the distance between two nodes v_*i*_^*R*^ and v_*j*_ ^*R*^ is less than 5 Å*A*, an edge e_*ij*_ ^*R*^ is established, and the edge length serves as the edge feature. If the distance between a protein node v_*i*_^*R*^ and a ligand node v_*i*_^*L*^ is less than 8 Å*A*, directed edges e_*i*,*i*_^*RL*^ and e_*i*,*i*_^*LR*^ are created, where e_*i*,*i*_^*RL*^ denotes the direction from v_*i*_^*R*^ to v_*i*_^*L*^, and e_*i*,*i*_^*LR*^ represents the direction from v_*i*_^*L*^ to v_*i*_^*R*^. This multilateral architecture facilitates the integration of node and edge information within the protein-ligand graph. To provide a more comprehensive description of the relative positions between protein-ligand atoms, in addition

## Short Article Title 3

to considering the interatomic distance, we introduce a novel orientation feature. Specifically, we introduce a dihedral angle *φ* (formed by the geometric center of the pose, v_*i*_^*L*^, v_*i*_^*R*^, and *α*-C), as well as two angles *θ*_1_ and *θ*_2_, as shown in Figure 1B. In Figure 1B, C_*α*_, R, L and P represent the C_*α*_ of the residue, a certain atom within the residue, a certain atom within the ligand and the geometric center of the ligand, respectively. *θ*_1_ and *θ*_2_ are the angles formed between the C*α*-R edge and the L-R edge, as well as P-L edge and R-L edge, respectively. Finally, sin(*φ*/2), cos(*θ*_1_/2), cos(*θ*_2_/2), and distance are utilized as features for the interaction edge. Table 1 summarizes all the node and edge features within this heterogeneous graph.

**Fig. 1.**
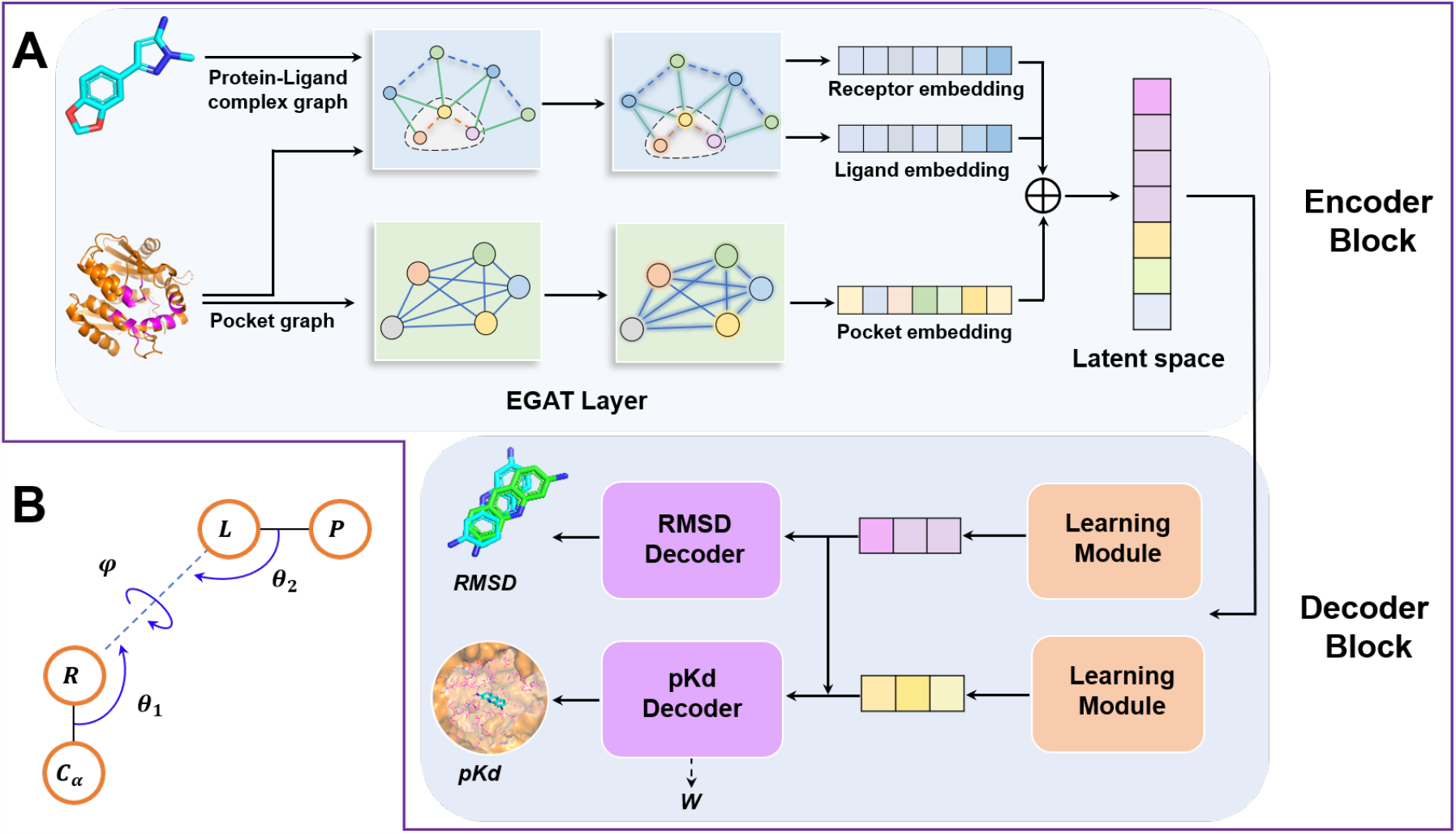
Overview of IGModel. A, Illustration of predicting RMSD and pKd from the protein-ligand complex structure. B, The orientation information on the relative positions of protein-ligand atoms. C_*α*_, R, L and P represent the C_*α*_ of the residue, a certain atom within the residue, a certain atom within the ligand, and the geometric center of the ligand, respectively. The *θ*_1_ and *θ*_2_ are the angles between C_*α*_-R and L-R, and between P-L and R-L, respectively. The *Ψ* denotes the dihedral angle formed by C_*α*_, R, L and P.

**Table 1.**
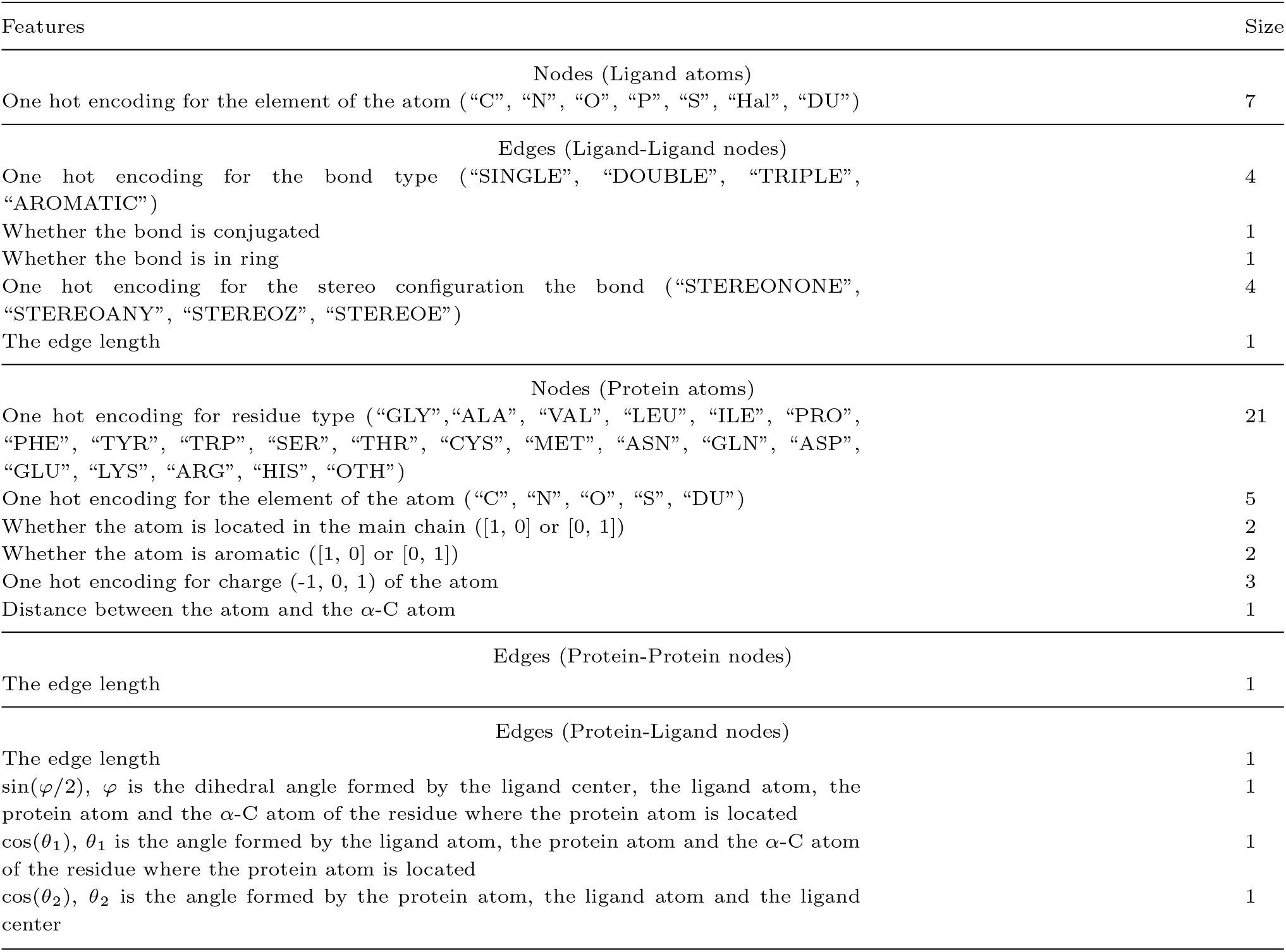
Node and Edge Features employed for protein-ligand interaction graph.

As we all know, the physicochemical environment of the binding pocket is critical for ligand binding[59, 60]. However, macroscopic descriptions of binding pockets cannot be effectively conveyed solely through atomic-level interaction graphs. To address this limitation, we construct an undirected graph G^*p*^=(V^*p*^, E^*p*^) to describe the residue states at the binding pocket. Similar to RTMScore, we defined the binding pocket as the residues within 8Å*A* around the co-crystallized ligand. Each node v_*i*_^*p*^*∈*V^*p*^ represents a residue within the binding pocket, and an edge e_*ij*_ ^*p*^ is established when the minimum distance between two residues is less than 10 Å*A*. Node features primarily encompass the residue type, distance distribution between internal atoms, and position relative to the pocket center, while edge features represent the distances between nodes in the main chain atoms. Table 2 summarizes the detailed node and edge features of the binding pocket graph.

**Table 2.** Node and Edge Features Employed for protein pocket graph.

| Features | Size |
| --- | --- |
| Nodes |  |
| One hot encoding for residue type (“GLY”, “ALA”, “VAL”, “LEU”, “ILE”, “PRO”, “PHE”, “TYR”, “TRP”, “SER”, “THR”, “CYS”, “MET”, “ASN”, “GLN”, “ASP”, “GLU”, “LYS”, “ARG”, “HIS”, “OTH”) | 21 |
| Max distance between any atom and α-C atom | 1 |
| Distance between atoms named C and N | 1 |
| Distance between α-C atom and the pocket center | 1 |
| Max and Min distance between any atom and the pocket center | 2 |
| $\sin(\phi/2)$ and $\sin(\psi/2)$, where $\phi$ and $\psi$ are main chain dihedral angles | 2 |
| $\sin(\chi_i/2)$, where $\chi_i$ ($i = 1, 2, 3, 4$ and $5$) is the side chain dihedral angle. If a dihedral angle does not exist in the residue, it is set to -2. | 5 |
| Edges |  |
| Distance between the α-C atoms of two residues | 1 |
| Distance between the main-chain carboxyl O atoms of two residues | 1 |
| Distance between the main-chain N atoms of two residues | 1 |
| Distance between the main-chain carboxyl C atoms of two residues | 1 |
| Distance between the centers of two residues | 1 |
| Max and min distance between two residues | 2 |

### Architecture

IGModel mainly contains three parts: the interaction feature encoding module, the RMSD decoding module and the pKd decoding module.

The encoding part of IGModel comprises two branches, dedicated to handing the binding pocket graph and the proteinligand interaction graph, with each branch containing two EdgeGAT layers. When the binding pocket graph is input into the encoder, its node features h_*pock*_ and edge features f_*pock*_ are updated as shown in Eq.1. The updated node features h^*′*^_*pock*_ is converted into a vector of length 1024 to represent the binding pocket embedding.

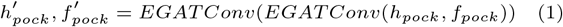

For the protein-ligand interaction graph, the first round of message passing is shown in Eq.2-5. After the first round of message passing, the nodes of protein and ligand atoms have been updated. The updated nodes of the protein and the ligand subgraphs, h^*′*^_*rec*_ and h^*′*^_*lig*_, are as shown in Eq.6-7. After two rounds of updates, the node features of the protein and the ligand are transformed into 1024-dimensional vectors, which serves as embeddings containing information about the protein and ligand atoms. Combining these two embeddings with the binding pocket embedding yields the final latent space describing the protein-ligand interaction.

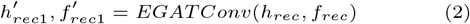

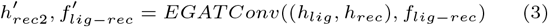

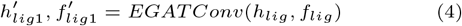

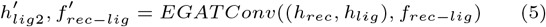

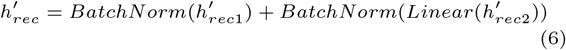

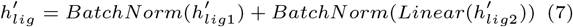

The two learning modules in the decoding part consist of a gMLP[61] layer and two linear layers. After passing through these two learning modules, the latent space is converted into two 128-dimensional vectors, namely V_*RMSD*_ and V_*pkd*_. In order to make the predicted pKd perceive the change of the RMSD of the ligand pose, we map V_*RMSD*_ to a new vector V^*′*^_*pkd*_ on the space of V_*pkd*_, which is integrated with V_*pkd*_ for decoding the pKd.

The training data consists of experimentally determined structures and virtual conformations generated by molecular docking. However, only the binding affinity of native complexes are available. Here, we assume that the binding strength of the binding pose against the target is inversely proportional to its RMSD relative to the native conformation, while the native protein-ligand conformation has the greatest binding affinity. In this work, we aim to learn the relationship between the RMSD of the docking pose and the binding strength through DL. In detail, the model also predicted a decay factor *W* (as shown in Figure 1A and Eq.8) with a value range of 0-1 in the pKd decoding part to describe the magnitude of the reduction of pKd with RMSD. Through *W* and the previous assumptions, the binding strength (pKd_*label*_) of the docking pose to the receptor can be deduced, this will serve as the label for pKd (Eq.9).

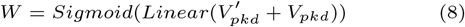

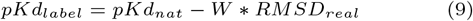

In Eq.8, pKd_*nat*_ is binding affinity between the native conformation of the ligand and the receptor, and RMSD_*real*_ is the real RMSD of the docking pose. When RMSD_*real*_ is close to 0, pKd_*label*_ will be close to pKd_*nat*_.

The loss function during training is defined as follows:

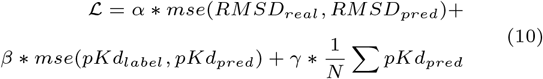

where a, b and c are the weights when summing up the components within the loss function. The last term in the loss function is used to constrain the convergence direction of pKd, thus improving the stability of pKd prediction.

## Results

### Evaluation on CASF-2016 benchmark

Distance information between protein and ligand atoms is crucial for describing protein-ligand interaction[25, 35, 21]. However, in this work, we have also incorporated relative positional information between protein and ligand atoms to enhance the characterization of interaction. As illustrated in Figure 2, we introduced two angles, *θ*_1_ and *θ*_2_, as well as a dihedral angle *ϕ*, which are obviously rotation invariant. Graph representation based on distance and direction will more comprehensively restore the geometric information of protein-ligand atoms.

**Fig. 2.**
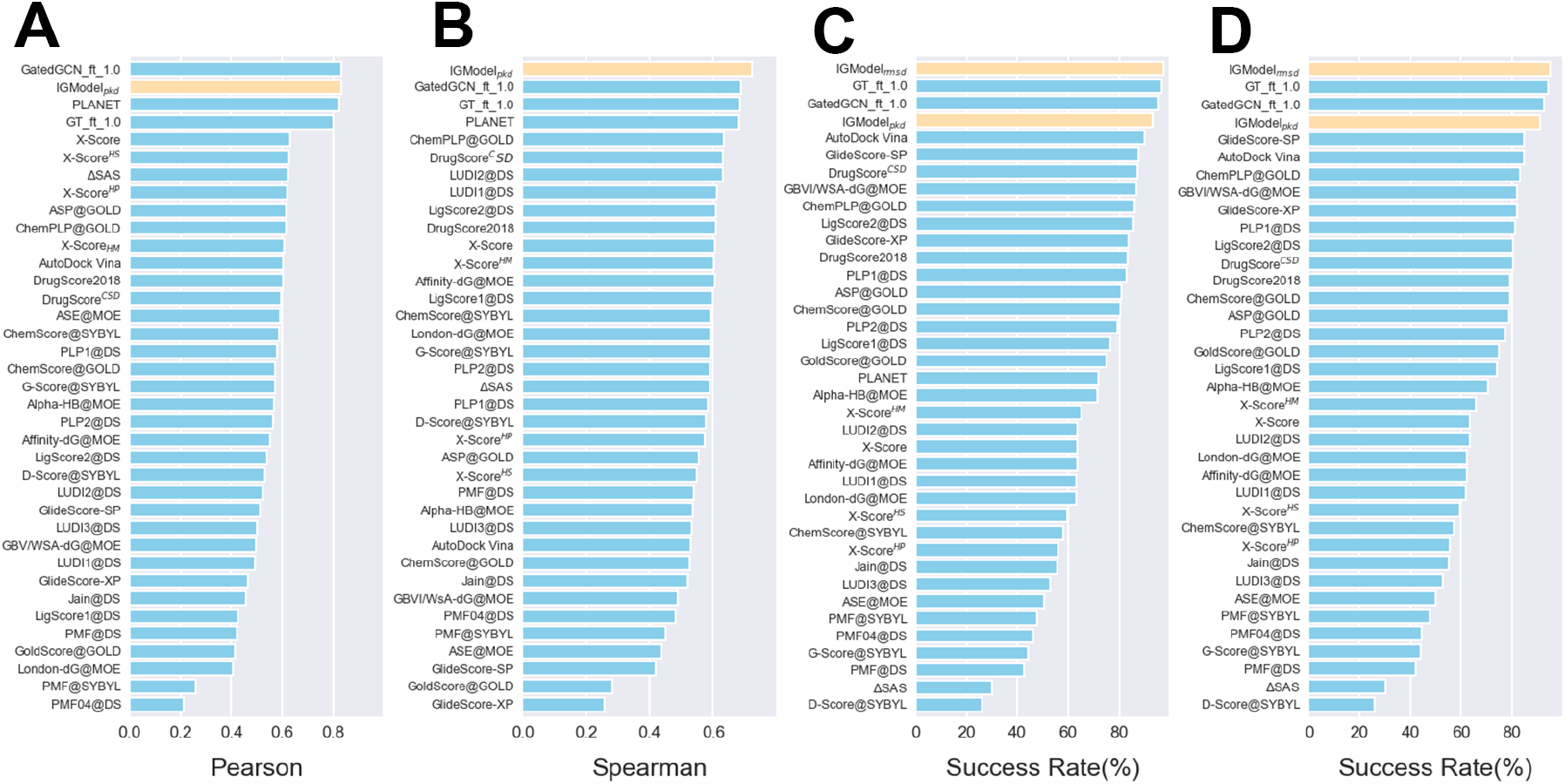
Comparison of the scoring power, ranking power and docking power in the CASF-2016 benchmark with other SFs. The CASF-2016 benchmark is compared with other traditional SFs reported in refxx, as well as some recently reported DL-based models. **A**. Scoring power measures the correlation between the scores of the model and experimental affinity, and the evaluation metric is the Pearson correlation coefficient. **B**. Ranking power evaluates the ability of a SF to rank the known ligands for a certain target, and its evaluation metric is the Spearman correlation coefficient. **C** and **D** show the top 1 success rate when the crystal structures are included and excluded from the test set, respectively.

The performance of the SF is generally evaluated from four aspects: scoring power, ranking power, docking power and screening power as defined in CASF-2016 benchmark[28]. Previous research has shown that some SFs with strong docking power or screening power often have poor scoring power and ranking power, such as DeepRMSD+Vina[39] and RTMScore[41]. In other words, training a model that can be applicable to various tasks with balanced performance is quite challenging[43]. Despite our model primarily focusing on identifying near-native conformation of the ligand (docking power), what is surprising is that it also has a relatively balanced performance in other tasks (Figure 2 and Table 3). Firstly, for docking power, the top 1 success rate of IGModel_*rmsd*_ with native poses included in the test set is 97.5%, and the value remains as high as 95.3% when the native poses are excluded. At the same time, IGModel_*pkd*_ can also achieve the higher top 1 success rates compared to most models, which are 93.0% and 90.0% when crystal structures are included and excluded in the test set, respectively. In addition, screening power refers to the ability of the SFs to identify the true binders to a specific receptor among a large library of compounds, which is measured by two indicators: the first one is the success rate of identifying the highest-affinity binder among the 1%, 5% or 10% top-ranked ligands over all 57 target proteins in the test set; the second indicator is enrichment factor (EF) that is calculated by the average percentage of the true binders among the 1%, 5% or 10% top-ranked candidates across all 57 targets. IGModel achieved a Top 1% success rate of 66.7%, which is comparable to RTMScore[41] but slightly lower than GT ft 1.0[43]. However, the EF achieved by IGModel_*pkd*_ is only 19.8, which is significantly lower than RTMScore and GT ft 1.0, but still higher than most predictors, such as DeepDock[40]. In general, IGModel exhibits a relatively balanced performance across various metrics based on CASF-2016 benchmark. The ablation experiments of IGModel on the validation set and CASF-2016 docking power are shown in Support Information part 3. Interestingly, for scoring power, IGModel_*pkd*_ achieved a Person correlation coefficient (PCC) of 0.831, which is very close to the 0.834 and 0.824 achieved by GatedCGN ft 1.0[43] and PLANET[42] repectively. The scatter plots of IGModel_*pkd*_ on the validation set and CASF-2016 core set are shown in Support Information part 2. Lastly, for ranking power test, IGModel_*pkd*_ achieved a Spearman correlation coefficient (SCC) of 0.723, which is higher than the 0.686 achieved by GatedCGN ft 1.0.

**Table 3.** The docking power and screening power of several representative SFs on the CASF-2016 benchmark.

| Scoring function | docking power |  | screening power |  |
| --- | --- | --- | --- | --- |
|  | top 1 success rate (w/o native poses) | top 1 success rate (with native poses) | top 1% success rate | enrichment factor |
| AutoDock Vina[28] | 0.846 | 0.902 | 0.298 | 7.70 |
| ChemPLP@GOLD[28] | 0.832 | 0.860 | 0.351 | 11.91 |
| GlideScore-SP[28] | 0.846 | 0.877 | 0.368 | 11.44 |
| $\Delta_{Vina}RF_{20}$[28] | 0.849 | 0.891 | 0.421 (0.456) | 11.73 (12.36) |
| $\Delta_{Vina}XGB$[62] | | 0.920 | 0.368 | 13.14 |
| $\Delta_{LinF9}XGB$[45] | | 0.867 | 0.404 | 12.61 |
| OnionNet-SFCT+Vina[44] |  | 0.937 | 0.421 | 15.50 |
| DeepBSP[38] | 0.872 | 0.885 |  |  |
| DeepDock[40] |  | 0.870 | 0.439 | 16.41 |
| RTMScore[41] | 0.934 | 0.973 | 0.667 | 28.0 |
| GT_ft_1.0[43] | 0.940 | 0.966 | 0.719 | 28.12 |
| GatedGCN_ft_1.0[43] | 0.926 | 0.954 | 0.661 | 23.54 |
| IGModel<sub>pkd</sub> | 0.909 | 0.933 | 0.667 | 19.40 |
| IGModel<sub>rmsd</sub> | 0.951 | 0.975 |  |  |

Note: The results of SFs other than IGModel are cited from the reference.

### Evaluation on the redocking and cross-docking test sets

Currently, in most molecular docking applications, the protein is treated as a rigid molecule, which deviated from the actual protein-ligand binding behavior observed in the real world, since the binding pocket could accommodate different compounds with flexible side-chains and sometimes also adjusted backbones[63]. Cross-docking refers to redocking a certain ligand to a non-cognate receptor[52]. Redocking and cross-docking are two ways for evaluating molecular docking, which refers to redocking a certain ligand to a cognate receptor and docking a ligand to a non-cognate receptor in the original pocket, respectively. To comprehensively assess the docking power of IGModel, various redocking and cross-docking benchmarks were adopted.

The first test set we assessed is PDBbind-CrossDocked-Core, with all receptors and ligands derived from the PDBbind v2016 core set. Each ligand was extracted from 285 proteinligand crystal structures, and then was redocked into the original protein or other four proteins belonging to the same target cluster by three docking softwares: Surflex-Dock, Glide SP and AutoDock Vina. IGModel was tested in these three groups of poses and compared with other SFs, and the results are shown in Table 4 and Figure 3. For cross-docking, the top 1 success rates on poses generated by IGModel_*rmsd*_ with Surflex, Glide and Vina are 0.662, 0.595 and 0.594 respectively, which is slightly better than GT ft 1.0 and GatedGCN ft 1.0, and significantly ahead of most predictors. Meanwhile, IGModel_*pkd*_ is still able to perform well that is comparable with GatedGCN ft 1.0, though it is slight worse than IGModel_*rmsd*_. For redocking tasks, the top 1 success rates of IGModel_*rmsd*_ and IGModel_*pkd*_ on poses generated by Surflex, Glide and Vina are 0.854 and 0.850, 0.786 and 0.779 as well as 0.761 and 0.754 respectively, which demonstrate a significant advantage over other SFs.

**Table 4.** Docking power (Top 1 success rate) of IGModel and other SFs on the PDBbind-CrossDocked-Core set.

| SFs | Surflex |  | Glide SP |  | Vina |  |
| --- | --- | --- | --- | --- | --- | --- |
|  | Redocking | Cross-docking | Redocking | Cross-docking | Redocking | Cross-docking |
| AD4 | 0.702 | 0.498 | 0.603 | 0.498 | 0.551 | 0.437 |
| Vina | 0.691 | 0.505 | 0.606 | 0.505 | 0.540 | 0.380 |
| Vinardo | 0.677 | 0.477 | 0.628 | 0.477 | 0.558 | 0.391 |
| X-Score | 0.663 | 0.475 | 0.582 | 0.475 | 0.512 | 0.401 |
| Pafnucy | 0.512 | 0.319 | 0.422 | 0.319 | 0.211 | 0.165 |
| Glide SP | 0.730 | 0.547 | 0.645 | 0.547 | 0.502 | 0.376 |
| Glide XP | 0.726 | 0.525 | 0.610 | 0.525 | 0.470 | 0.366 |
| GT_ft_1.0 | 0.815 | 0.636 | 0.743 | 0.585 | 0.660 | 0.590 |
| GatedGCN_ft_1.0 | 0.822 | 0.627 | 0.719 | 0.575 | 0.674 | 0.581 |
| IGModel<sub>pkd</sub> | 0.850 | 0.626 | 0.779 | 0.577 | 0.754 | 0.565 |
| IGModel<sub>rmsd</sub> | 0.854 | 0.662 | 0.786 | 0.595 | 0.761 | 0.594 |

Note: The results of SFs other than IGModel are cited from reference[43].

**Fig. 3.**
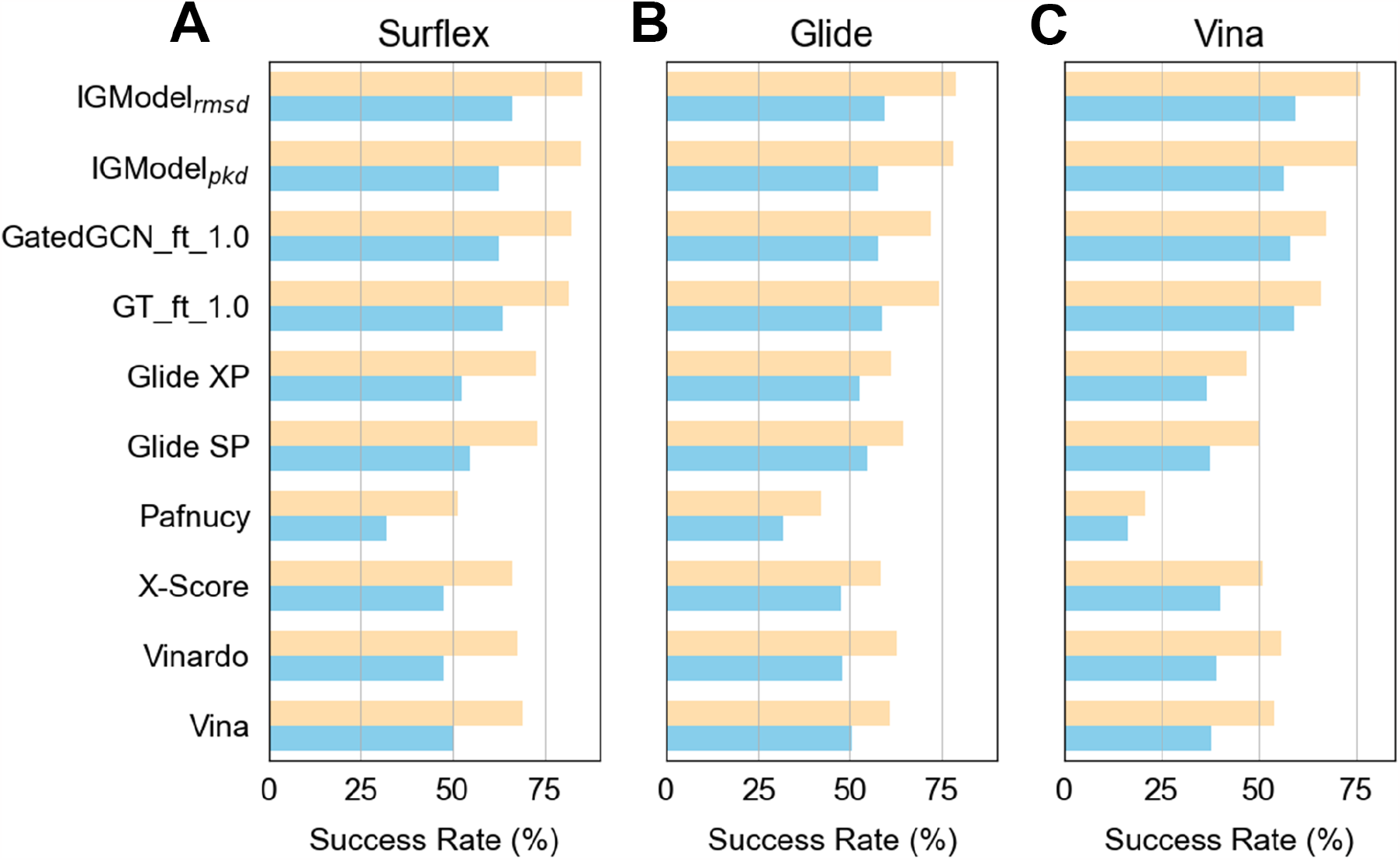
Docking power of IGModel and other SFs on PDBbind-CrossDocked-Core set. **A, B** and **C** show the top 1 success rate of SFs on poses generated by Surflex, GLide and AutoDock Vina, respectively.

Another cross-docking test set used in this study is DISCO, which contains 4399 crystal protein-ligand complexes across 95 protein targets[52]. These targets are sourced from DUD-E[64], thus covering a wide range of protein families. The poses of ligands are generated by AutoDock Vina[55], with 20 poses generated for each protein-ligand pair by default. In order to be consistent with our previous research, each specific target-ligand was treated as a single case when calculating the top1 success rate, which is different from the integrated idea when tested on PDBbind-CrossDocked-Core set. The performance of IGModel compared to several representative SFs on DISCO set is shown in Figure 4. It can be clearly seen that IGModel_*rmsd*_ is ahead of other baseline SFs, while IGModel_*pkd*_ performs comparably to GatedGCN ft 1.0. The excellent performance of IGModel in redocking and cross-docking tasks proves its significant practical application value.

**Fig. 4.**
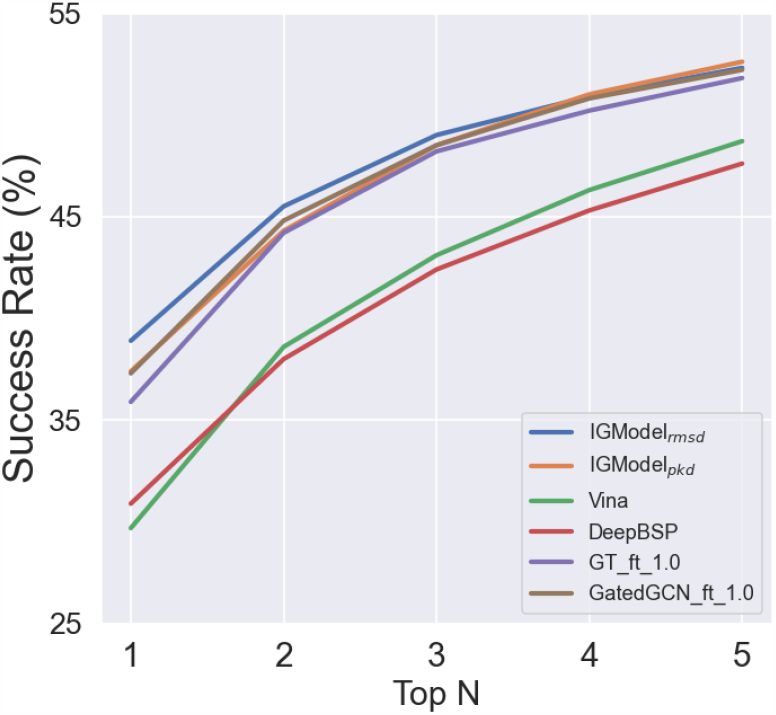
The top N success rate of IGModel and other baseline models on DISCO set.

### Generalization Assessment of IGModel

In this paper, we assessed the generalization ability of the IGModel from two additional perspectives. Firstly, most of SFs do not rigorously eliminate redundancy in the training set, which may lead to the results of SFs on the test set being much better than in actual scenarios. Therefore, we introduced a unbiased test set (called unbias-v2019 set in this work) that we previously proposed, which contains the protein-ligand pairs with low similarity to those in PDBbind database v2019, and the native conformation of the ligands were excluded[46]. The similarity is defined as the product of protein sequence similarity (calculated by NW-align) and the Tanimoto similarity of the ligand Morgan fingerprints calculatied by rdkit[57]. Secondly, when the native conformation of the target is unknown, predicting the binding pose of the ligand becomes even more challenging. Computation methods such as AlphaFold2[53] provide solutions rapidly obtaining high-precision protein structures. Then, on the basis of these predicted protein structures, molecular docking was applied to generate poses of the ligands. In our previously study, CASF-2016 and the unbias-v2019 were used to construct the test sets for the AlphaFold2 version, namely CASF-2016-AF2 and unbias-2019-AF2, respectively[46].

We assessed the docking power of SFs on the three datasets mentioned above. Table 5 shows the Pearson correlation coefficient (PCC) and Spearman correlation coefficient (SCC) between the scores of SFs and the true RMSD. It can be clearly seen that, compared to other baseline SFs, IGModel exhibits remarkably high accuracy. Especially on the unbias-v2019-AF2 set, IGModel can still perform robustly. The Top1

**Table 5.** Pearson and Spearman correlation coefficient of IGModel and other SFs tested on the CASF2016-AF2, unbias-v2019 and unbias-v2019-AF2 datasets.

| SFs | CASF2016-AF2 |  | unbias-v2019 |  | unbias-v2019-AF2 |  |
| --- | --- | --- | --- | --- | --- | --- |
|  | PCC | SCC | PCC | SCC | PCC | SCC |
| Vina | 0.035 | 0.071 | 0.364 | 0.356 | 0.209 | 0.192 |
| DeepBSP | 0.401 | 0.375 | 0.543 | 0.507 | 0.451 | 0.418 |
| DeepRMSD | 0.463 | 0.430 | 0.285 | 0.246 | 0.261 | 0.247 |
| DeepRMSD+Vina | 0.248 | 0.290 | 0.405 | 0.362 | 0.299 | 0.287 |
| GT_ft_1.0 | 0.503 | 0.455 |  |  |  |  |
| GatedGCN_ft_1.0 | 0.560 | 0.524 |  |  |  |  |
| zPoseScore | 0.659 | 0.593 | 0.604 | 0.535 | 0.554 | 0.507 |
| IGModel<sub>pkd</sub> | 0.736 | 0.649 | 0.611 | 0.522 | 0.580 | 0.519 |
| IGModel<sub>rmsd</sub> | 0.751 | 0.669 | 0.633 | 0.542 | 0.609 | 0.552 |

Note: The results of SFs other than IGModel are cited from reference[46].

success rate achieved by SFs with 2Å*A* and 3Å*A* as cutoffs is shown in Table 6, and what is exciting is that IGModel still significantly outperforms other SFs. This effectively verifies the generalization ability and robustness of IGModel in different situations. For CASF2016-AF2 and unbias-v2019-AF2, two datasets based on the structures predicted by AlphaFold2, since the poses generated by molecular docking have larger RMSDs relative to the native poses, the overall top1 success rate is lower.

**Table 6.** Top1 success rate of IGModel and other SFs tested on the CASF2016-AF2, unbias-v2019 and unbias-v2019-AF2 datasets.

| SFs | CASF2016-AF2 |  | unbias-v2019 |  | unbias-v2019-AF2 |  |
| --- | --- | --- | --- | --- | --- | --- |
|  | 2Å | 3Å | 2Å | 3Å | 2Å | 3Å |
| Vina | 0.127 | 0.208 | 0.453 | 0.526 | 0.130 | 0.219 |
| DeepBSP | 0.180 | 0.324 | 0.453 | 0.584 | 0.169 | 0.254 |
| DeepRMSD | 0.269 | 0.425 | 0.210 | 0.324 | 0.089 | 0.185 |
| DeepRMSD+Vina | 0.261 | 0.416 | 0.441 | 0.534 | 0.144 | 0.253 |
| GT_ft_1.0 | 0.273 | 0.448 |  |  |  |  |
| GatedGCN_ft_1.0 | 0.292 | 0.465 |  |  |  |  |
| zPoseScore | 0.339 | 0.506 | 0.599 | 0.700 | 0.185 | 0.274 |
| IGModel<sub>pkd</sub> | 0.338 | 0.558 | 0.562 | 0.690 | 0.227 | 0.318 |
| IGModel<sub>rmsd</sub> | 0.364 | 0.571 | 0.587 | 0.690 | 0.227 | 0.296 |

Note: The results of SFs other than IGModel are cited from reference[46].

## Discussion

Different SFs were developed with various training data for either pKd or RMSD predictions, and a single score (either a traditional SF or a ML/DL-based SF) may perfectly address the docking pose quality[44] regarding the pose selection and virtual screening[28]. For pose selection, RMSD reflects the difference between the docking pose and the native conformation (thus solving the “how it binds” problem), but cannot represent the binding strength with the target protein; while for virtual screening, pKd represents the binding strength between the molecule and the target (regarding the “how strong it binds” problem), but cannot reflect the difference between the docking pose and the native pose [65].

Early ML-based or DL-based SFs usually directly predict protein-ligand binding affinity (pKd) given the native proteinligand complex structures [25, 34, 35, 36]. However, testing shows that the scores of such models are difficult to distinguish correct binding poses generated by docking tools [28], and their accuracies on screening tasks are neither satisfied[37]. Later, researchers began to directly predict the RMSD of the docking poses (such as Gnina [66], DeepRMSD[39] and DeepBSP[38]), or use scores from other mathematical spaces (such as DeepDock[40], RTMScore[41] and GenScore[43]) but trained with docking poses. The distance likelihood potential generated by DeepDock, RTMScore and Genscore is difficult to intuitively describe RMSD and the binding strength and could not provide an explicit and physical meaningful predictions for computational chemists. Whereas, RMSD corrected pKd is also used as training target to construct ML models, it could provide direct estimate for both pose binding pattern and molecule binding strength with a single score[62, 45].

To this end, we try to answer the two questions (RMSD prediction for “how it binds” and pKd prediction for “how strong it binds”) within one integrated framework, where we characterize the protein-ligand interaction through two geometric graph modules: the pocket graph module and the protein-ligand atomic graph module, and then apply EdgeGAT layers[50] to encode the interaction features. Tests under different scenarios have shown that IGModel is good at predicting the RMSD of the docking poses for both redocking, cross-docking and AF2-based docking tasks, indicating that it has a excellent generalization ability and robustness for pose selection (Tables 3, 4 and 5).

In order to more clearly display the protein-ligand interaction potential energy surface encoded by the graph neural network, we visualized the ligand embedding output by the last EdgeGAT layer in the protein-ligand graph branch and the overall complex embedding. First, we cluster the latent space and the ligand embedding of the samples in the validation set through principal component analysis (PCA), and then color them according to real RMSD, predicted RMSD, pKd_*label*_ and predicted pKd respectively. The distributions of RMSD and pKd on the latent space are shown in Figure 5 A-D. It can be clearly found that as the RMSD and pKd change, obvious layering appears on the pattern. Similar trends can also be observed in ligand embedding pattern (Figure 5 E-H). However, there is a significant phenomenon that the latent space image has a larger coverage area compared to the ligand embedding, which means that the latent space is better able to distinguish the differences between different clusters. This may be attributed to the integration of protein pocket information within the latent space, thus enhancing the representation of protein-ligand interactions.

**Fig. 5.**
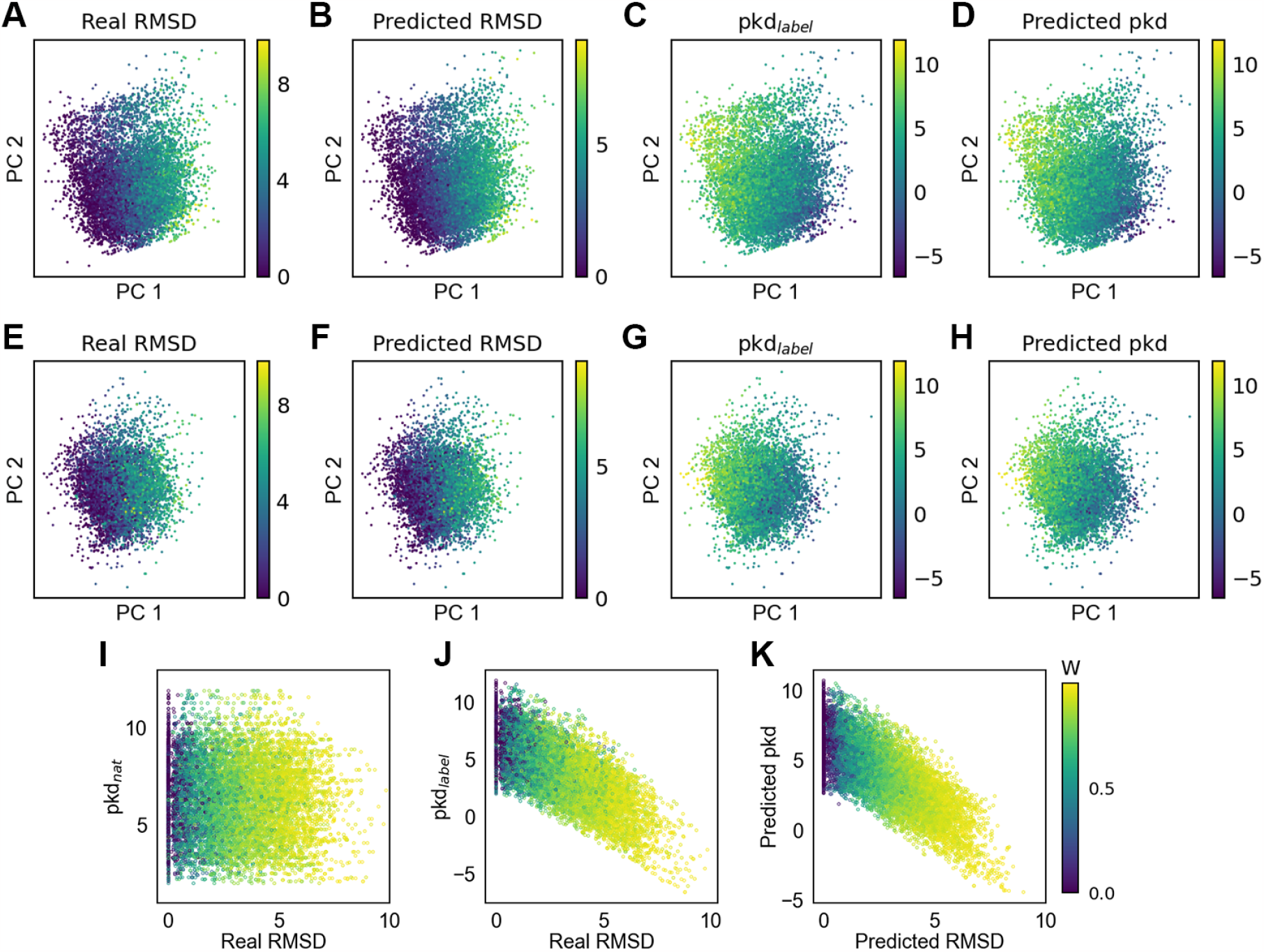
Visualization of the complex embedding, ligand embedding and the decay factor W. A-D (E-H) show the visualization of the complex embedding (ligand embedding) of samples in the validation set after PCA clustering, colored according to real RMSD, predicted RMSD, pKd_*label*_ and predicted pKd, respectively. I-K present visualization of the decay weight of binding strength with respect to RMSD variations, organized according to pKd_*nat*_-real RMSD, pKd_*label*_-real RMSD and predicted pKd-Predicted RMSD, respectively. Where pKd_*nat*_ refers to the binding affinity of the native protein-ligand complex.

IGModel, as the first deep learning model capable of simultaneously predicting the RMSD of ligand docking poses relative to the native conformation and the binding strength to the target, is founded on the assumption of the negative correlation between binding strength and RMSDs of docking poses. However, the decay weight of binding strength with respect to the RMSDs of docking poses is unknown. Previously, researchers have empirically set functions for the variation of binding strength with RMSD[62, 45], but such manual interventions may introduce systematic errors. Therefore, we hope that the model can obtain the corresponding decay weights based on different protein-ligand complexes (as shown in Figure 1). We display the decay weights according to pKd_*nat*_-real RMSD (Figure 5I), pKd_*label*_-real RMSD (Figure 5J) and predicted pKd-predicted RMSD (Figure 5K) respectively. One clear trend is that as the RMSD increases, the decay weight *W* also increases. This aligns with the initial assumption that poses closer to the native conformation have the stronger binding strength.

Next we also explore the ability of the model to learning physical protein-ligand interactions. It is assumed that protein-ligand binding is majorly driven by non-bonded interactions [67], solvation effects [68] and entropic effects [69]. For non-bonded interactions, in particular, short-range interactions such as hydrophobic interactions, cation-*π* interactions, salt bridges, hydrogen bonds and *π*-*π* stacking usually provide favorable binding free energies for the protein-ligand complex [70, 71, 67]. However, most current ML/DL-based SFs fail to explicitly or implicitly emphasize or highlight the direct non-bonded interactions between the protein and the ligand.

Atoms located in different residue side chains, and even those at different positions within the same residue, often exhibit distinct physicochemical properties. Therefore, comprehensively considering multiple features such as residue type, element type, main chain/side chain, polar/non-polar and aromaticity, effectively assigns physiologically relevant identifiers to protein atoms. This makes it possible for IGModel to capture key non-bonded interactions such as hydrogen bonding (Figure 6). We extracted the attention values generated by IGModel for each protein-ligand atom pair in the protein-ligand complexes from CASF-2016 docking poses to examine whether the physical interactions are highlighted or have been paid enough “attentions”. The importance of a protein atom is defined as the sum of attention values of the protein-ligand interaction edges in which this protein atom participates. For example, if a protein atom forms edges with five ligand atoms, the importance of the protein atom is represented by the sum of these five attention values. Subsequently, the importance of all protein atom types was counted, with top 20 ranked types displayed in Figure 6 A. We can clearly find that seven of top eight most important protein atoms are polar atoms, which implies the significant role of polar interactions in protein-ligand binding[70] for binding strength and binding specificity. In addition, certain non-polar atoms, such as ILE-CD1 and PHE-CZ, also play a crucial role. This is because alpha carbon atoms and aromatic rings typically participate in the hydrophobic interactions, while it is the most frequently occurring interaction type in protein-ligand binding. Figure 6B shows that the protein atoms involved in hydrogen bonding have higher importance values, which shows that IGModel is capable of recognizing hydrogen bonds and assigning them greater attention. In Figure 6C, the periphery of the benzene ring in phenylalanine that is close to the benzene ring of the small molecule also has a high importance value, which indicates that IGModel has the ability to detect *π*-*π* stacking.

**Fig. 6.**
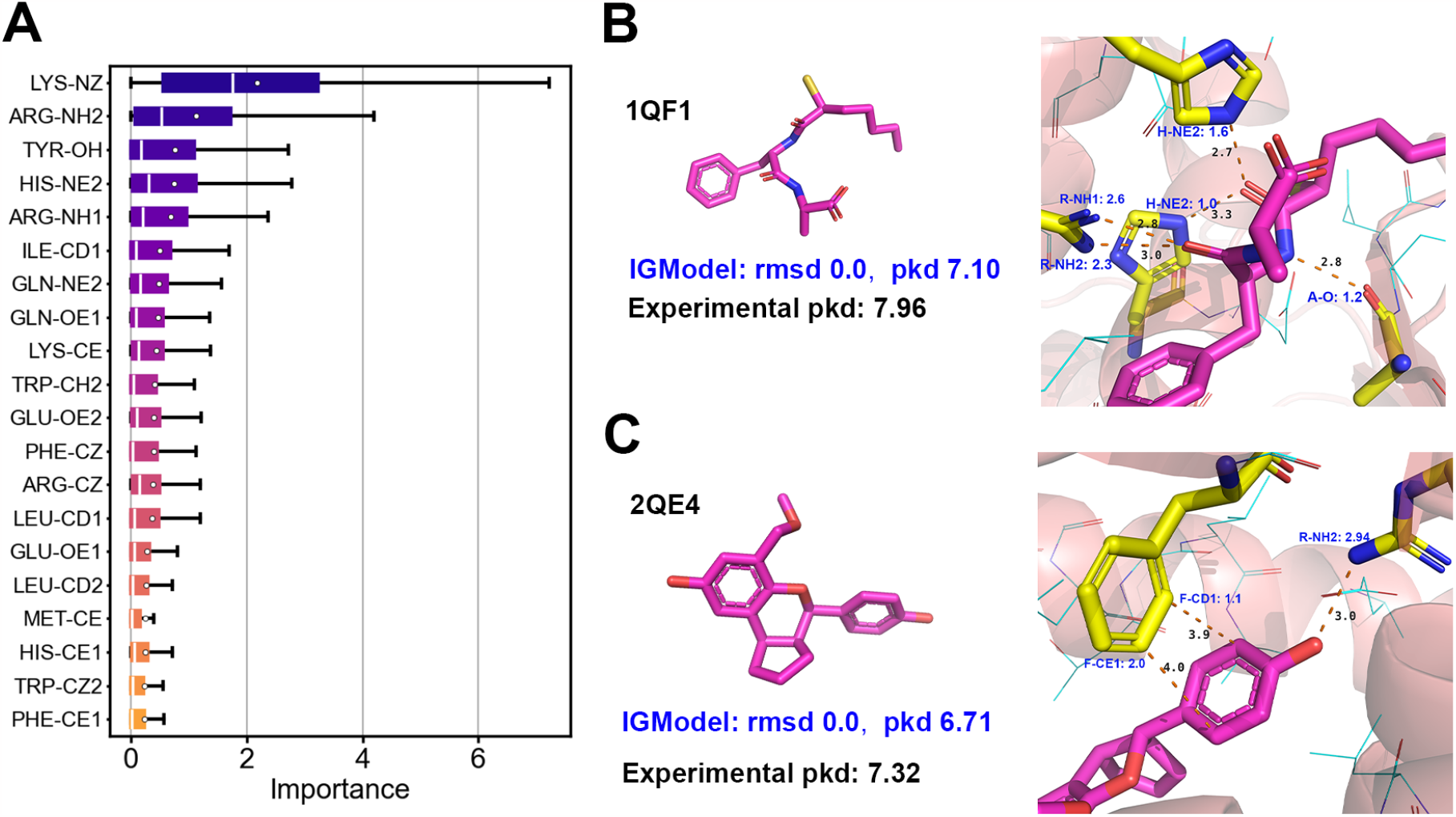
Explainability and case study. A. Ranking of importance of protein atoms at the binding pocket (only the top 20 protein atom types are shown). B and C show two cases (1QF1 and 2QE4) where IGModel identifies key interactions, including hydrogen bonding and *π*-*π* stacking. In the two pictures located on the right, the distances between key atoms are shown (black), and the names and importance values of key atoms in the protein are colored in blue.

### Conclusion

In this work, we propose a new scoring framework called IGModel for protein-ligand interaction prediction, which can simultaneously predicts the RMSD of the ligand binding pose and the binding strength with the target. IGModel applies EdgeGAT layer to encode the two input graphs into the latent space characterizing the protein-ligand interaction, and then decodes the latent space into RMSD and pKd through two decoders respectively. The results show that it achieves SOTA performance in almost all docking power test sets. Although IGModel aims to provide a more comprehensive quality assessment for docking poses, it still performs well on the CASF-2016 scoring power and ranking power test, which is comparable or even better to the other models. For screening power, out model is also ahead of most baseline SFs. Furthermore, IGModel is also evaluated on the more challenging unbiased set unbias-v2019 and data set containing target structure predicted by AlphaFold2, proving its strong generalization capabilities. We also visualized the latent space encoded by IGModel, providing an intuitive representation of the energy space describing the RMSD and binding strength of the docking pose. Through case studies, it was observed that IGModel is capable of identifying critical interactions, such as hydrogen bonding and *π*-*π* stacking.

It is undeniable that one SF is difficult to perform perfectly in all tasks, but IGModel can achieve a relatively balanced performance. The most importance is that IGModel is a new framework for predicting protein-ligand interactions, which breaks the tradition that the SFs only output a single score, and ensures that the output values have intuitive physical meanings. We believe that this framework is valuable for molecular docking and even the modification and optimization of lead compounds in drug design. In summary, our research proposes a new paradigm for the design of SFs in the future, along withe new challenges.

## Author contributions statement

Z.W. designed research, performed research, analyzed data and wrote the paper; S.W., Y.L., J.G. and Y.W. analyzed data; Y.M. analyzed data and wrote the paper; L.Z. and W.L. designed research, analyzed data and wrote the paper.

## Acknowledgments

This work is partly supported by the Singapore Ministry of Education (MOE), tier 1 grants RG97/22 (M.Y.). This work was partly supported by the Key Research and Development Project of Guangdong Province under grant no. 2021B0101310002, National Science Foundation of China under grant no. 62272449, the Shenzhen Basic Research Fund under grant no RCYX20200714114734194, KQTD20200820113106007, ZDSYS20220422103800001. We would also like to thank the funding support by the Youth Innovation Promotion Association(Y2021101), CAS to Yanjie Wei.

## Author Name

This is sample author biography text. The values provided in the optional argument are meant for sample purposes. There is no need to include the width and height of an image in the optional argument for live articles. This is sample author biography text this is sample author biography text this is sample author biography text this is

sample author biography text this is sample author biography text this is sample author biography text this is sample author biography text this is sample author biography text.

## Author Name

This is sample author biography text this is sample author biography text this is sample author biography text this is sample author biography text this is sample author biography text this is sample author biography text this is sample author biography text this is sample author biography text.

## Supporting Materials

### Part 1. Details of docking poses in the training set

In this work, docking poses were generated by AutoDock Vina and ledock, where the search space is a 15Å*A*\*15Å*A*\*15Å*A* cubic, the maximum number generated is set to 10, and other parameters are used by default. The RMSD distribution of docking poses and the pKd distribution of the native conformations corresponding to these docking poses are shown in Figure S1.

**Fig. S1.**
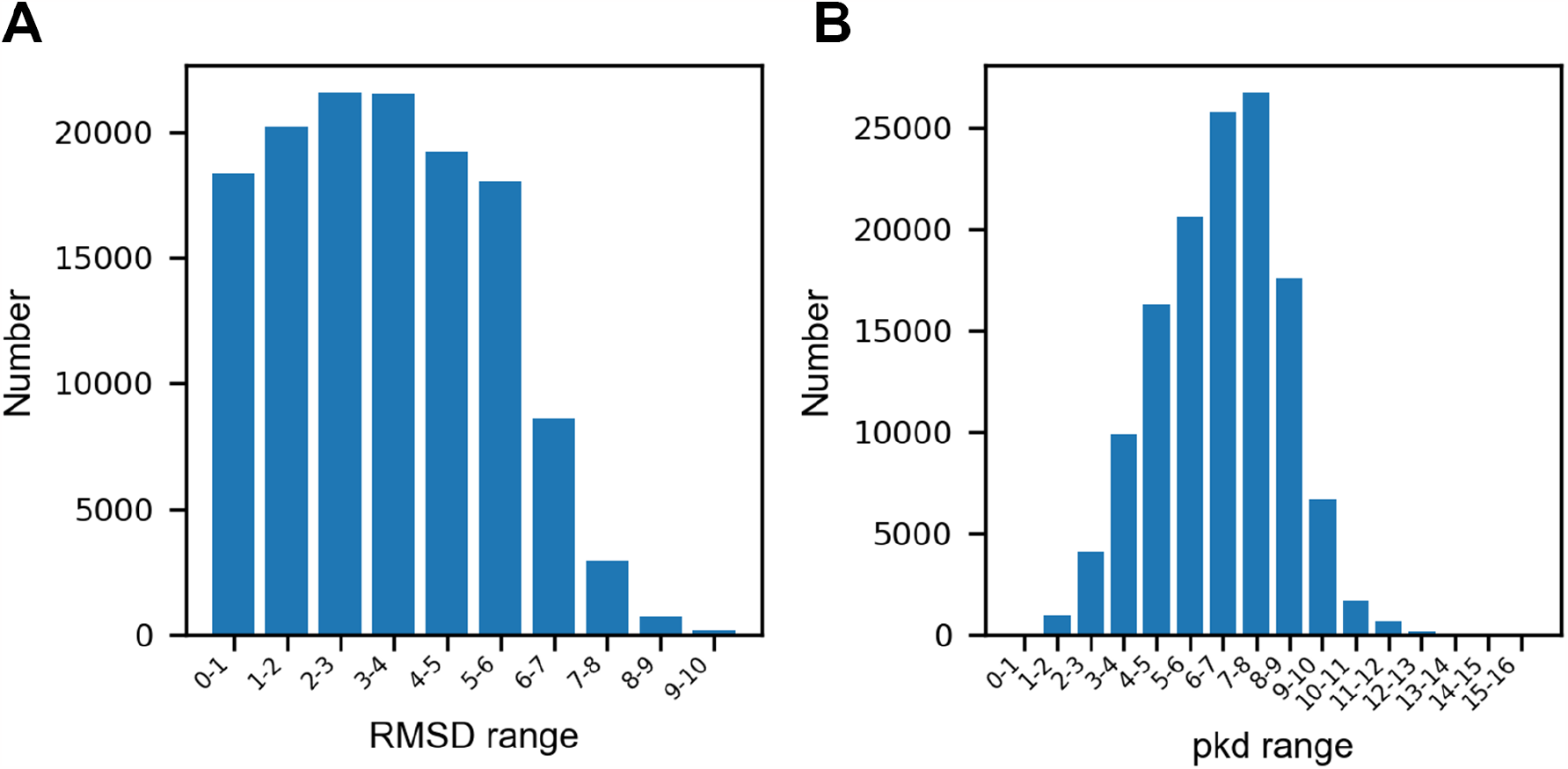
The RMSD (A) and pKd (B) distributions of docking poses in the training set.

### Part 2. Performance of IGModel and some representative scoring functions in CASF-2016 benchmark

The Pearson correlation coefficient (PCC) and root-mean-squared error (RMSE) achieved by IGModel_*pkd*_ on CASF-2016 core set are 0.831 and 1.254, respectivily, as shown in Figure S2A. Figure S2B shows the results of IGModel_*pkd*_ on the validation set, including crystal structures and decoys. The performance of IGModel_*pkd*_ and several recently published representative models on CASF-2016 scoring power in Table S1.

**Table S1.** The scoring power of IGModel_*pkd*_ and other representative scoring functions in CASF-2016 benchmark.

| Year | SFs | PCC | RMSE | Training set |
| --- | --- | --- | --- | --- |
| 2023 | IGModel<sub>pkd</sub> | 0.831 | 1.254 | PDBbind v2019 (general set) |
| 2023 | GatedGCN_ft_1.0 | 0.834 |  | PDBbind v2020 (general set) |
| 2023 | GT_ft_1.0 | 0.802 |  | PDBbind v2020 (general set) |
| 2023 | PLANET | 0.824 | 1.247 | PDBbind v2020 (general set) |
| 2022 | RTMScore | 0.455 |  | PDBbind v2020 (general set) |
| 2022 | MPNN | 0.813 | 1.511 | PDBbind v2016 (general set) |

### Part 3. The ablation study of IGModel

As shown in Table S2, when removing Angle 1, Angle 2, dihedral angle formed between the protein atoms and the ligand atoms in the protein-ligand interaction graph, or removing the pocket graph, the performance of IGModel shows varying degrees of decrease.

**Fig. S2.**
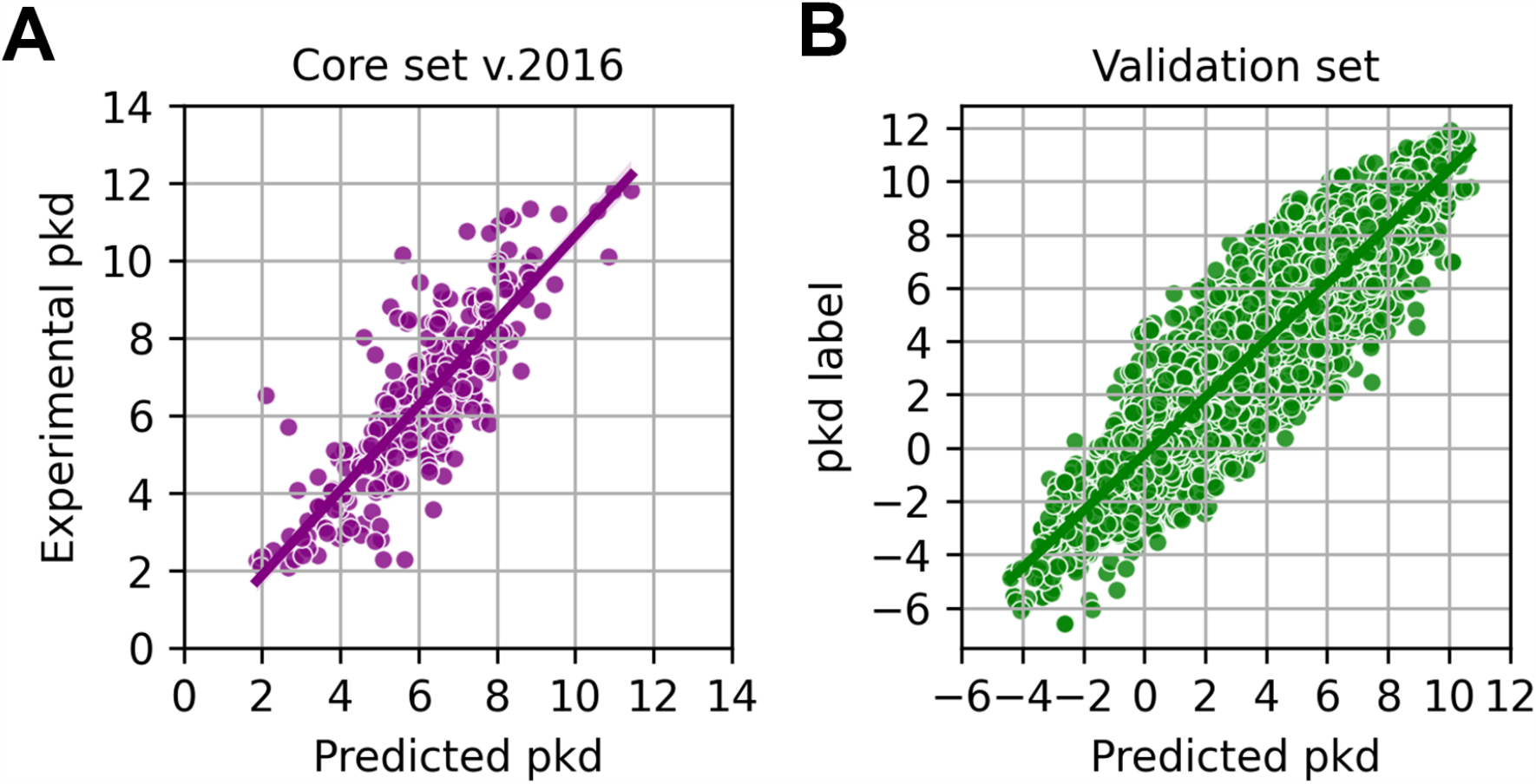
A. The correlation between the pKd predicted by IGModel_*pkd*_ and the experimental pKd for 285 native protein-ligand complexes in the CASF-2016 core set. B. The scatter plot of pKd predicted by IGModel_*pkd*_ against the pKd labels on the validation set.

**Table S2.**
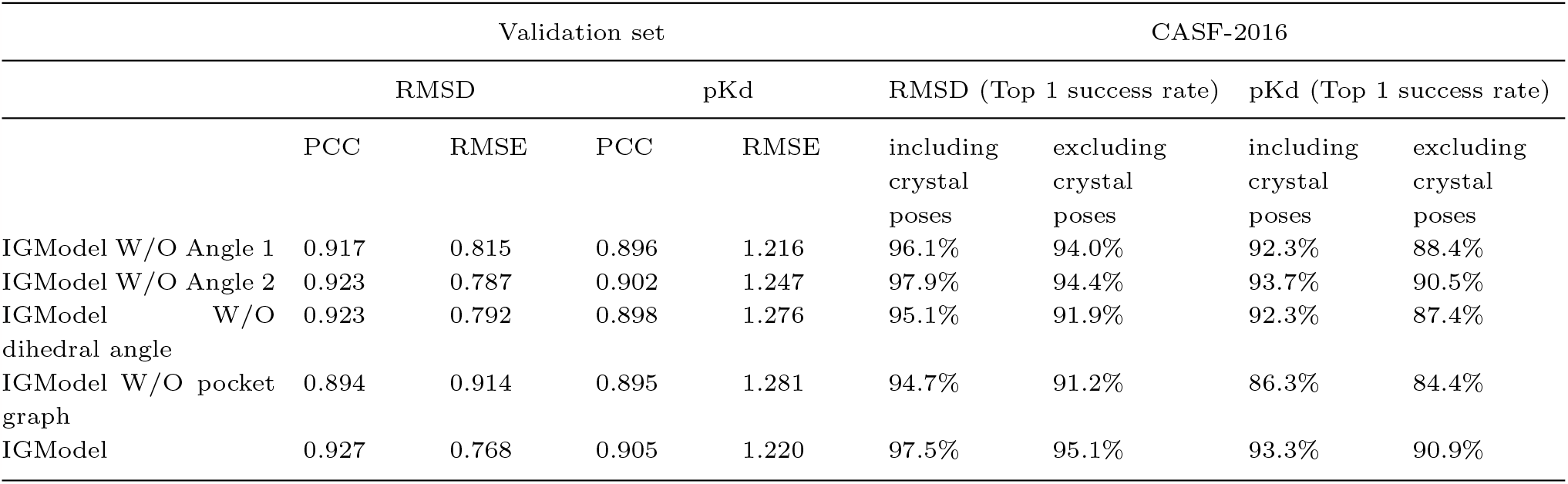
The ablation experiment of IGModel on the validation set and CASF-2016 docking power.

|  | Validation set |  |  |  | CASF-2016 |  |  |  |
| --- | --- | --- | --- | --- | --- | --- | --- | --- |
|  | RMSD |  | pKd |  | RMSD (Top 1 success rate) |  | pKd (Top 1 success rate) |  |
|  | PCC | RMSE | PCC | RMSE | including crystal poses | excluding crystal poses | including crystal poses | excluding crystal poses |
| IGModel W/O Angle 1 | 0.917 | 0.815 | 0.896 | 1.216 | 96.1% | 94.0% | 92.3% | 88.4% |
| IGModel W/O Angle 2 | 0.923 | 0.787 | 0.902 | 1.247 | 97.9% | 94.4% | 93.7% | 90.5% |
| IGModel W/O dihedral angle | 0.923 | 0.792 | 0.898 | 1.276 | 95.1% | 91.9% | 92.3% | 87.4% |
| IGModel W/O pocket graph | 0.894 | 0.914 | 0.895 | 1.281 | 94.7% | 91.2% | 86.3% | 84.4% |
| IGModel | 0.927 | 0.768 | 0.905 | 1.220 | 97.5% | 95.1% | 93.3% | 90.9% |

## References

1. Amy C Anderson. The process of structure-based drug design. Chemistry & biology, 10(9):787–797, 2003.

2. Luca Pinzi and Giulio Rastelli. Molecular docking: shifting paradigms in drug discovery. International journal of molecular sciences, 20(18):4331, 2019.

3. Sam Z Grinter and Xiaoqin Zou. Challenges, applications, and recent advances of protein-ligand docking in structure-based drug design. Molecules, 19(7):10150–10176, 2014.

4. Kathleen E Prosser, Ryjul W Stokes, and Seth M Cohen. Evaluation of 3-dimensionality in approved and experimental drug space. ACS medicinal chemistry letters, 11(6):1292–1298, 2020.

5. Stephani Joy Y Macalino, Vijayakumar Gosu, Sunhye Hong, and Sun Choi. Role of computer-aided drug design in modern drug discovery. Archives of pharmacal research, 38:1686–1701, 2015.

6. Riccardo Guareschi, Iva Lukac, Ian H Gilbert, and Fabio Zuccotto. Sophosqm: Accurate binding affinity prediction in compound optimization. ACS omega, 8(17):15083–15098, 2023.

7. Debleena Paul, Gaurav Sanap, Snehal Shenoy, Dnyaneshwar Kalyane, Kiran Kalia, and Rakesh K Tekade. Artificial intelligence in drug discovery and development. Drug discovery today, 26(1):80, 2021.

8. Thomas J Lane. Protein structure prediction has reached the single-structure frontier. Nature Methods, 20(2):170–173, 2023.

9. Rocco Meli, Garrett M Morris, and Philip C Biggin. Scoring functions for protein-ligand binding affinity prediction using structure-based deep learning: A review. Frontiers in bioinformatics, 2:57, 2022.

10. María JR Yunta. Docking and ligand binding affinity: uses and pitfalls. Am. J. Model. Optim, 4(3):74–114, 2016.

11. Xing Du, Yi Li, Yuan-Ling Xia, Shi-Meng Ai, Jing Liang, Peng Sang, Xing-Lai Ji, and Shu-Qun Liu. Insights into protein–ligand interactions: mechanisms, models, and methods. International journal of molecular sciences, 17(2):144, 2016.

12. Leonardo G Ferreira, Ricardo N Dos Santos, Glaucius Oliva, and Adriano D Andricopulo. Molecular docking and structure-based drug design strategies. Molecules, 20(7):13384–13421, 2015.

13. Pedro HM Torres, Ana CR Sodero, Paula Jofily, and Floriano P Silva-Jr. Key topics in molecular docking for drug design. International journal of molecular sciences, 20(18):4574, 2019.

14. Surovi Saikia and Manobjyoti Bordoloi. Molecular docking: challenges, advances and its use in drug discovery perspective. Current drug targets, 20(5):501–521, 2019.

15. Hilbert Yuen In Lam, Robbe Pincket, Hao Han, Xing Er Ong, Zechen Wang, Jamie Hinks, Yanjie Wei, Weifeng Li, Liangzhen Zheng, and Yuguang Mu. Application of variational graph encoders as an effective generalist algorithm in computer-aided drug design. Nature Machine Intelligence, 5(7):754–764, 2023.

16. Mohammad Hassan Baig, Khurshid Ahmad, Sudeep Roy, Jalaluddin Mohammad Ashraf, Mohd Adil, Mohammad Haris Siddiqui, Saif Khan, Mohammad Amjad Kamal, Ivo Provazník, and Inho Choi. Computer aided drug design: success and limitations. Current pharmaceutical design, 22(5):572–581, 2016.

17. Sofia D’Souza, KV Prema, and Seetharaman Balaji. Machine learning models for drug–target interactions: current knowledge and future directions. Drug Discovery Today, 25(4):748–756, 2020.

18. Jin Li, Ailing Fu, and Le Zhang. An overview of scoring functions used for protein–ligand interactions in molecular docking. Interdisciplinary Sciences: Computational Life Sciences, 11:320–328, 2019.

19. Francis E Agamah, Gaston K Mazandu, Radia Hassan, Christian D Bope, Nicholas E Thomford, Anita Ghansah, and Emile R Chimusa. Computational/in silico methods in drug target and lead prediction. Briefings in bioinformatics, 21(5):1663–1675, 2020.

20. Renee L DesJarlais, Robert P Sheridan, George L Seibel, J Scott Dixon, Irwin D Kuntz, and R Venkataraghavan. Using shape complementarity as an initial screen in designing ligands for a receptor binding site of known three-dimensional structure. Journal of medicinal chemistry, 31(4):722–729, 1988.

21. Sheng-You Huang, Sam Z Grinter, and Xiaoqin Zou. Scoring functions and their evaluation methods for protein– ligand docking: recent advances and future directions. Physical Chemistry Chemical Physics, 12(40):12899–12908, 2010.

22. Wei Deng, Curt Breneman, and Mark J Embrechts. Predicting protein-ligand binding affinities using novel geometrical descriptors and machine-learning methods. Journal of chemical information and computer sciences, 44(2):699–703, 2004.

23. Qurrat Ul Ain, Antoniya Aleksandrova, Florian D Roessler, and Pedro J Ballester. Machine-learning scoring functions to improve structure-based binding affinity prediction and virtual screening. Wiley Interdisciplinary Reviews: Computational Molecular Science, 5(6):405–424, 2015.

24. Antonio Lavecchia. Machine-learning approaches in drug discovery: methods and applications. Drug discovery today, 20(3):318–331, 2015.

25. Pedro J Ballester and John BO Mitchell. A machine learning approach to predicting protein–ligand binding affinity with applications to molecular docking. Bioinformatics, 26(9):1169–1175, 2010.

26. Jacob D Durrant and J Andrew McCammon. Nnscore: a neural-network-based scoring function for the characterization of protein-ligand complexes. Journal of chemical information and modeling, 50(10):1865–1871, 2010.

27. Duc Duy Nguyen and Guo-Wei Wei. Agl-score: algebraic graph learning score for protein–ligand binding scoring, ranking, docking, and screening. Journal of chemical information and modeling, 59(7):3291–3304, 2019.

28. Minyi Su, Qifan Yang, Yu Du, Guoqin Feng, Zhihai Liu, Yan Li, and Renxiao Wang. Comparative assessment of scoring functions: the casf-2016 update. Journal of chemical information and modeling, 59(2):895–913, 2018.

29. Hongming Chen, Ola Engkvist, Yinhai Wang, Marcus Olivecrona, and Thomas Blaschke. The rise of deep learning in drug discovery. Drug discovery today, 23(6):1241–1250, 2018.

30. Yankang Jing, Yuemin Bian, Ziheng Hu, Lirong Wang, and Xiang-Qun Sean Xie. Deep learning for drug design: an artificial intelligence paradigm for drug discovery in the big data era. The AAPS journal, 20:1–10, 2018.

31. Zhenyu Meng and Kelin Xia. Persistent spectral–based machine learning (perspect ml) for protein-ligand binding affinity prediction. Science advances, 7(19):eabc5329, 2021.

32. Joseph Gomes, Bharath Ramsundar, Evan N Feinberg, and Vijay S Pande. Atomic convolutional networks for predicting protein-ligand binding affinity. arXiv preprint arXiv:1703.10603, 2017.

33. José Jiménez, Miha Skalic, Gerard Martinez-Rosell, and Gianni De Fabritiis. K deep: protein–ligand absolute binding affinity prediction via 3d-convolutional neural networks. Journal of chemical information and modeling, 58(2):287–296, 2018.

34. Marta M Stepniewska-Dziubinska, Piotr Zielenkiewicz, and Pawel Siedlecki. Development and evaluation of a deep learning model for protein–ligand binding affinity prediction. Bioinformatics, 34(21):3666–3674, 2018.

35. Liangzhen Zheng, Jingrong Fan, and Yuguang Mu. Onionnet: a multiple-layer intermolecular-contact-based convolutional neural network for protein–ligand binding affinity prediction. ACS omega, 4(14):15956–15965, 2019.

36. Zechen Wang, Liangzhen Zheng, Yang Liu, Yuanyuan Qu, Yong-Qiang Li, Mingwen Zhao, Yuguang Mu, and Weifeng Li. Onionnet-2: a convolutional neural network model for predicting protein-ligand binding affinity based on residue-atom contacting shells. Frontiers in chemistry, 9:753002, 2021.

37. Chao Shen, Ye Hu, Zhe Wang, Xujun Zhang, Jinping Pang, Gaoang Wang, Haiyang Zhong, Lei Xu, Dongsheng Cao, and Tingjun Hou. Beware of the generic machine learning-based scoring functions in structure-based virtual screening. Briefings in Bioinformatics, 22(3):bbaa070, 2021.

38. Jingxiao Bao, Xiao He, and John ZH Zhang. Deepbsp—a machine learning method for accurate prediction of protein– ligand docking structures. Journal of chemical information and modeling, 61(5):2231–2240, 2021.

39. Zechen Wang, Liangzhen Zheng, Sheng Wang, Mingzhi Lin, Zhihao Wang, Adams Wai-Kin Kong, Yuguang Mu, Yanjie Wei, and Weifeng Li. A fully differentiable ligand pose optimization framework guided by deep learning and a traditional scoring function. Briefings in Bioinformatics, 24(1):bbac520, 2023.

40. Oscar Méndez-Lucio, Mazen Ahmad, Ehecatl Antonio del Rio-Chanona, and Jörg Kurt Wegner. A geometric deep learning approach to predict binding conformations of bioactive molecules. Nature Machine Intelligence, 3(12):1033–1039, 2021.

41. Chao Shen, Xujun Zhang, Yafeng Deng, Junbo Gao, Dong Wang, Lei Xu, Peichen Pan, Tingjun Hou, and Yu Kang. Boosting protein–ligand binding pose prediction and virtual screening based on residue–atom distance likelihood potential and graph transformer. Journal of Medicinal Chemistry, 65(15):10691–10706, 2022.

42. Xiangying Zhang, Haotian Gao, Haojie Wang, Zhihang Chen, Zhe Zhang, Xinchong Chen, Yan Li, Yifei Qi, and Renxiao Wang. Planet: A multi-objective graph neural network model for protein–ligand binding affinity prediction. Journal of Chemical Information and Modeling, 2023.

43. Chao Shen, Xujun Zhang, Chang-Yu Hsieh, Yafeng Deng, Dong Wang, Lei Xu, Jian Wu, Dan Li, Yu Kang, Tingjun Hou, et al. A generalized protein–ligand scoring framework with balanced scoring, docking, ranking and screening powers. Chemical Science, 14(30):8129–8146, 2023.

44. Liangzhen Zheng, Jintao Meng, Kai Jiang, Haidong Lan, Zechen Wang, Mingzhi Lin, Weifeng Li, Hongwei Guo, Yanjie Wei, and Yuguang Mu. Improving protein–ligand docking and screening accuracies by incorporating a scoring function correction term. Briefings in Bioinformatics, 23(3):bbac051, 2022.

45. Chao Yang and Yingkai Zhang. Delta machine learning to improve scoring-ranking-screening performances of protein– ligand scoring functions. Journal of Chemical Information and Modeling, 62(11):2696–2712, 2022.

46. Shen Tao, Liu Fuxu, Wang Zechen, Sun Jinyuan, Bu Yifan, Meng Jintao, Chen Weihua, Yao Keyi, Mu Yuguang, Li Weifeng, Zhao Guoping, Wang Sheng, Wei Yanjie, and Zheng Liangzhen. zposescore model for accurate and robust protein-ligand docking pose scoring in casp15. Proteins, 2023.

47. Ayan Chatterjee, Robin Walters, Zohair Shafi, Omair Shafi Ahmed, Michael Sebek, Deisy Gysi, Rose Yu, Tina Eliassi-Rad, Albert-Lászĺo Barabási, and Giulia Menichetti. Improving the generalizability of protein-ligand binding predictions with ai-bind. Nature Communications, 14(1):1989, 2023.

48. Huaipan Jiang, Jian Wang, Weilin Cong, Yihe Huang, Morteza Ramezani, Anup Sarma, Nikolay V Dokholyan, Mehrdad Mahdavi, and Mahmut T Kandemir. Predicting protein–ligand docking structure with graph neural network. Journal of chemical information and modeling, 62(12):2923–2932, 2022.

49. Zehong Zhang, Lifan Chen, Feisheng Zhong, Dingyan Wang, Jiaxin Jiang, Sulin Zhang, Hualiang Jiang, Mingyue Zheng, and Xutong Li. Graph neural network approaches for drug-target interactions. Current Opinion in Structural Biology, 73:102327, 2022.

50. Kamil Kamiński, Jan Ludwiczak, Maciej Jasinśki, Adriana Bukala, Rafal Madaj, Krzysztof Szczepaniak, and Stanis-law Dunin-Horkawicz. Rossmann-toolbox: a deep learning-based protocol for the prediction and design of cofactor specificity in rossmann fold proteins. Briefings in bioinformatics, 23(1):bbab371, 2022.

51. Chao Shen, Xueping Hu, Junbo Gao, Xujun Zhang, Haiyang Zhong, Zhe Wang, Lei Xu, Yu Kang, Dongsheng Cao, and Tingjun Hou. The impact of cross-docked poses on performance of machine learning classifier for protein–ligand binding pose prediction. Journal of cheminformatics, 13(1):1–18, 2021.

52. Shayne D Wierbowski, Bentley M Wingert, Jim Zheng, and Carlos J Camacho. Cross-docking benchmark for automated pose and ranking prediction of ligand binding. Protein Science, 29(1):298–305, 2020.

53. John Jumper, Richard Evans, Alexander Pritzel, Tim Green, Michael Figurnov, Olaf Ronneberger, Kathryn Tunyasuvunakool, Russ Bates, Augustin Žídek, Anna Potapenko, et al. Highly accurate protein structure prediction with alphafold. Nature, 596(7873):583–589, 2021.

54. Renxiao Wang, Xueliang Fang, Yipin Lu, and Shaomeng Wang. The pdbbind database: Collection of binding affinities for protein-ligand complexes with known three-dimensional structures. Journal of medicinal chemistry, 47(12):2977–2980, 2004.

55. Oleg Trott and Arthur J Olson. Autodock vina: improving the speed and accuracy of docking with a new scoring function, efficient optimization, and multithreading. Journal of computational chemistry, 31(2):455–461, 2010.

56. Ni Liu and Zhibin Xu. Using ledock as a docking tool for computational drug design. In IOP Conference Series: Earth and Environmental Science, volume 218, page 012143. IOP Publishing, 2019.

57. Greg Landrum et al. Rdkit: Open-source cheminformatics software. 2016.

58. Rocco Meli and Philip C Biggin. spyrmsd: symmetry-corrected rmsd calculations in python. Journal of Cheminformatics, 12(1):1–7, 2020.

59. Alfonso T García-Sosa. Hydration properties of ligands and drugs in protein binding sites: tightly-bound, bridging water molecules and their effects and consequences on molecular design strategies. Journal of chemical information and modeling, 53(6):1388–1405, 2013.

60. Francesca Spyrakis, Mostafa H Ahmed, Alexander S Bayden, Pietro Cozzini, Andrea Mozzarelli, and Glen E Kellogg. The roles of water in the protein matrix: a largely untapped resource for drug discovery. Journal of medicinal chemistry, 60(16):6781–6827, 2017.

61. Hanxiao Liu, Zihang Dai, David So, and Quoc V Le. Pay attention to mlps. Advances in Neural Information Processing Systems, 34:9204–9215, 2021.

62. Jianing Lu, Xuben Hou, Cheng Wang, and Yingkai Zhang. Incorporating explicit water molecules and ligand conformation stability in machine-learning scoring functions. Journal of chemical information and modeling, 59(11):4540–4549, 2019.

63. Jiyu Fan, Ailing Fu, and Le Zhang. Progress in molecular docking. Quantitative Biology, 7:83–89, 2019.

64. Michael M Mysinger, Michael Carchia, John J Irwin, and Brian K Shoichet. Directory of useful decoys, enhanced (dud-e): better ligands and decoys for better benchmarking. Journal of medicinal chemistry, 55(14):6582–6594, 2012.

65. David Ramírez and Julio Caballero. Is it reliable to take the molecular docking top scoring position as the best solution without considering available structural data? Molecules, 23(5):1038, 2018.

66. Andrew T McNutt, Paul Francoeur, Rishal Aggarwal, Tomohide Masuda, Rocco Meli, Matthew Ragoza, Jocelyn Sunseri, and David Ryan Koes. Gnina 1.0: molecular docking with deep learning. Journal of cheminformatics, 13(1):1–20, 2021.

67. Jiřrí Černý and Pavel Hobza. Non-covalent interactions in biomacromolecules. Physical Chemistry Chemical Physics, 9(39):5291–5303, 2007.

68. GL Amidon, RS Pearlman, and ST Anik. The solvent contribution to the free energy of protein-ligand interactions. Journal of Theoretical Biology, 77(1):161–170, 1979.

69. Lili Duan, Xiao Liu, and John ZH Zhang. Interaction entropy: A new paradigm for highly efficient and reliable computation of protein–ligand binding free energy. Journal of the American Chemical Society, 138(17):5722–5728, 2016.

70. Earl Frieden. Non-covalent interactions: Key to biological flexibility and specificity. Journal of chemical education, 52(12):754, 1975.

71. Katharina Wendler, Jens Thar, Stefan Zahn, and Barbara Kirchner. Estimating the hydrogen bond energy. The Journal of Physical Chemistry A, 114(35):9529–9536, 2010.

